## Supplementary figures and images for "Cardiac Fibroblasts regulate myocardium and coronary vasculature development via the collagen signaling pathway"

### Supplemental data 1

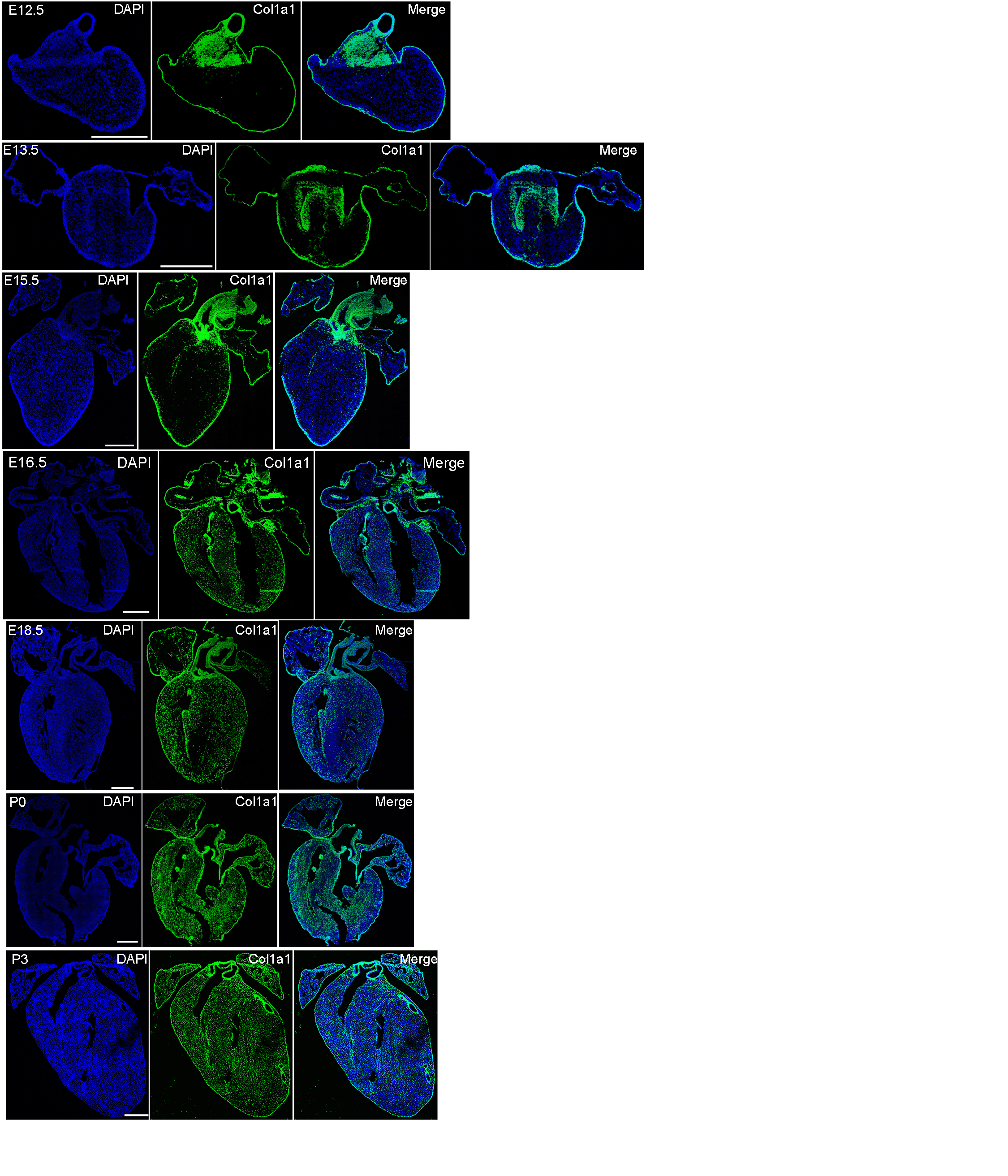

### Supplemental data 2

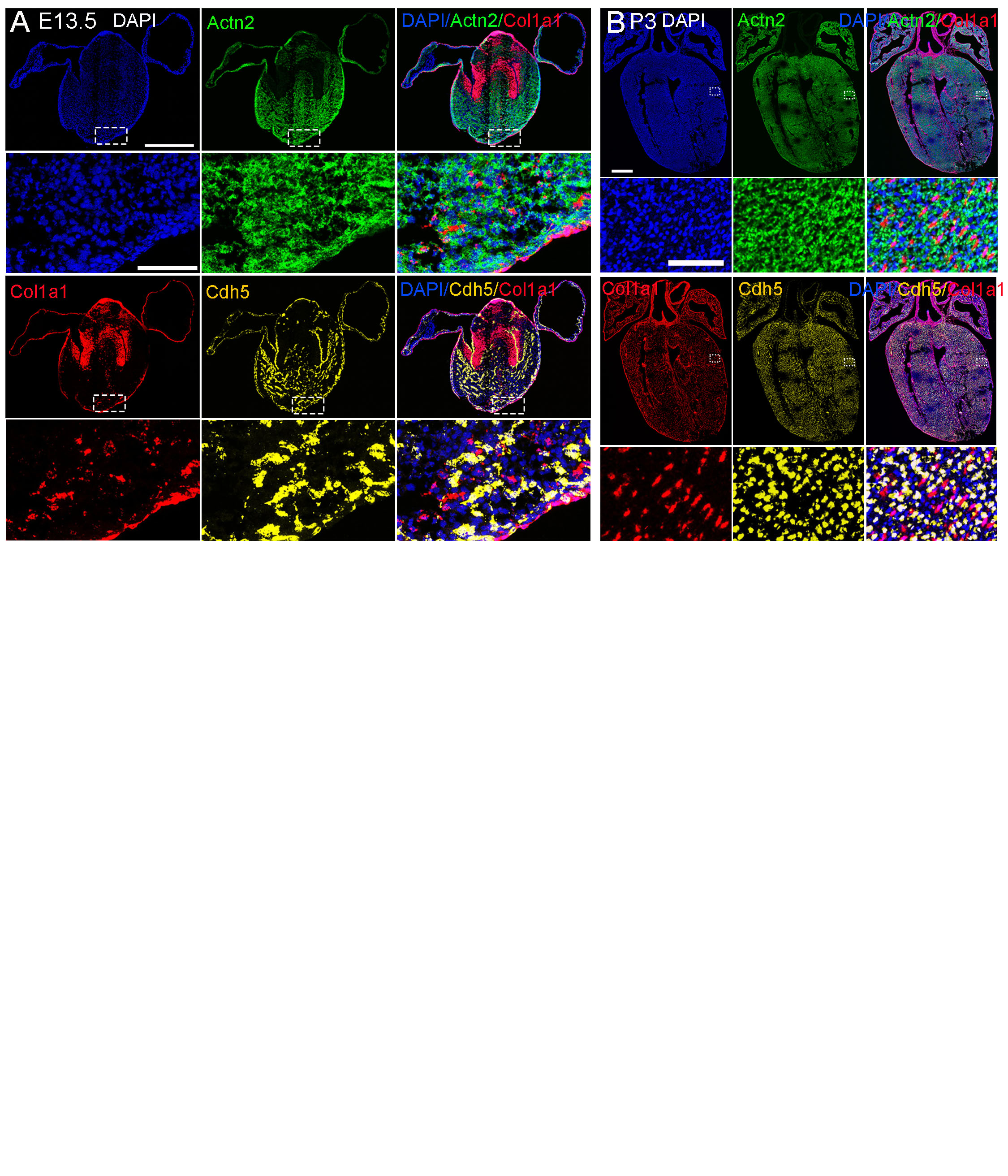

### Supplemental data 3

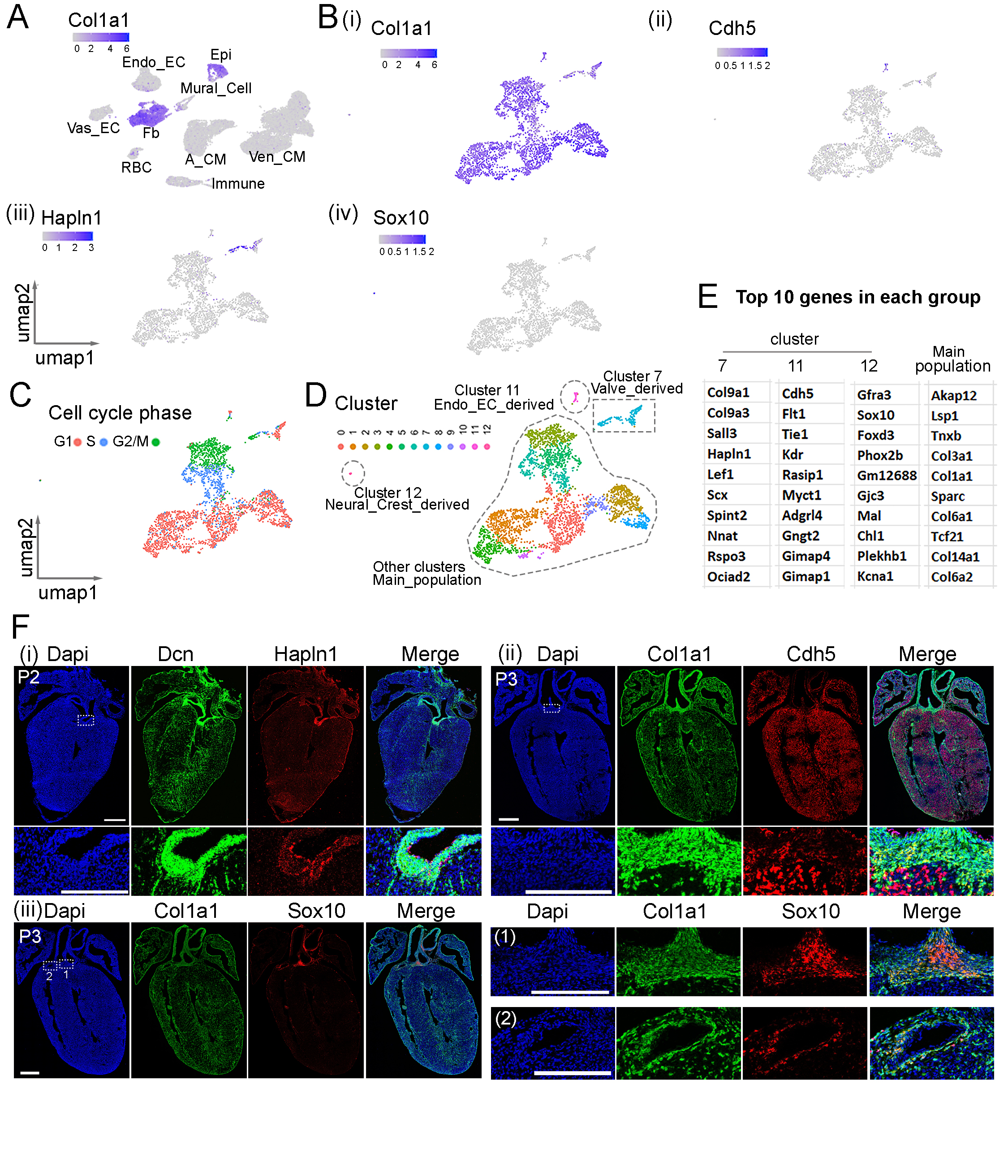

### Supplemental data 4

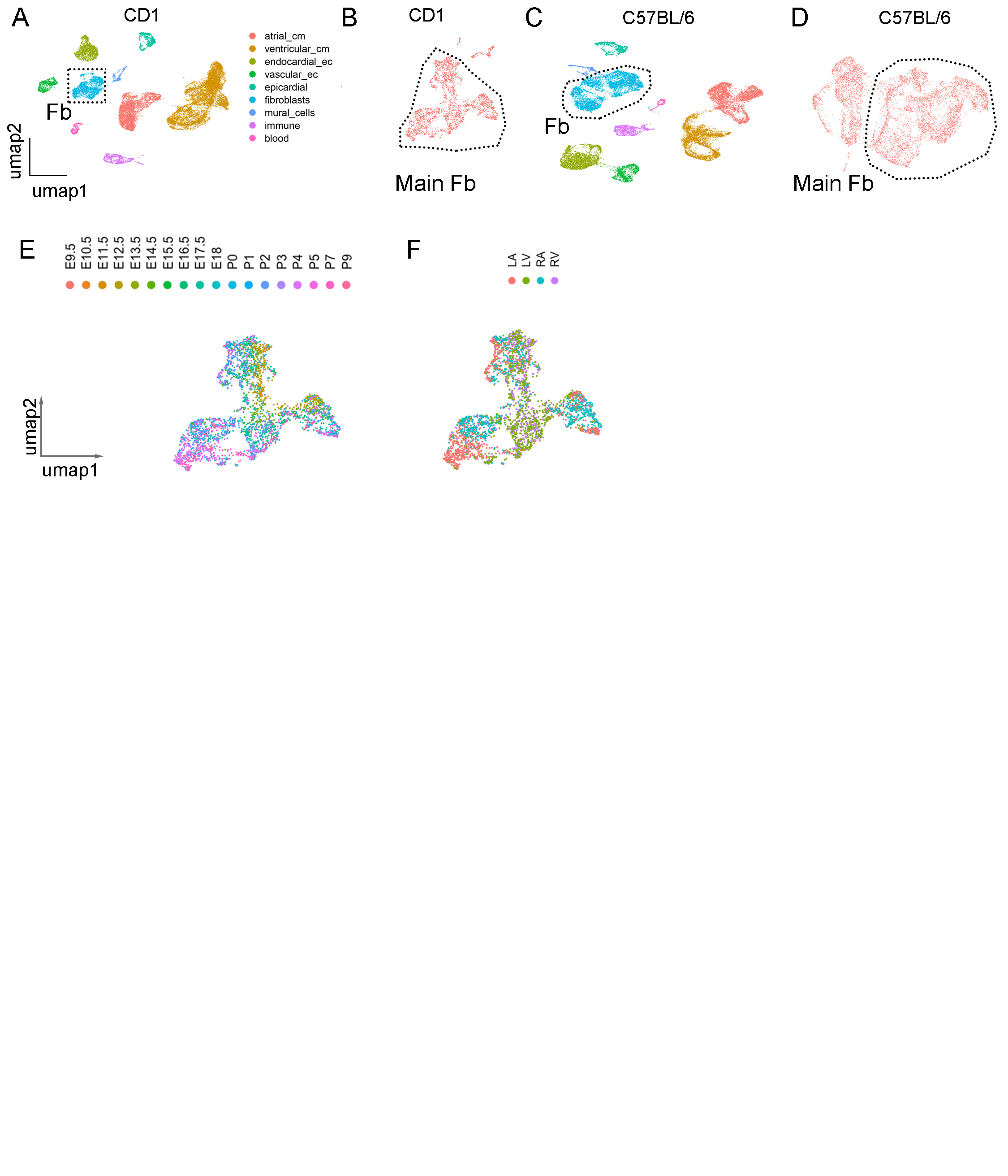

### Supplemental data 5

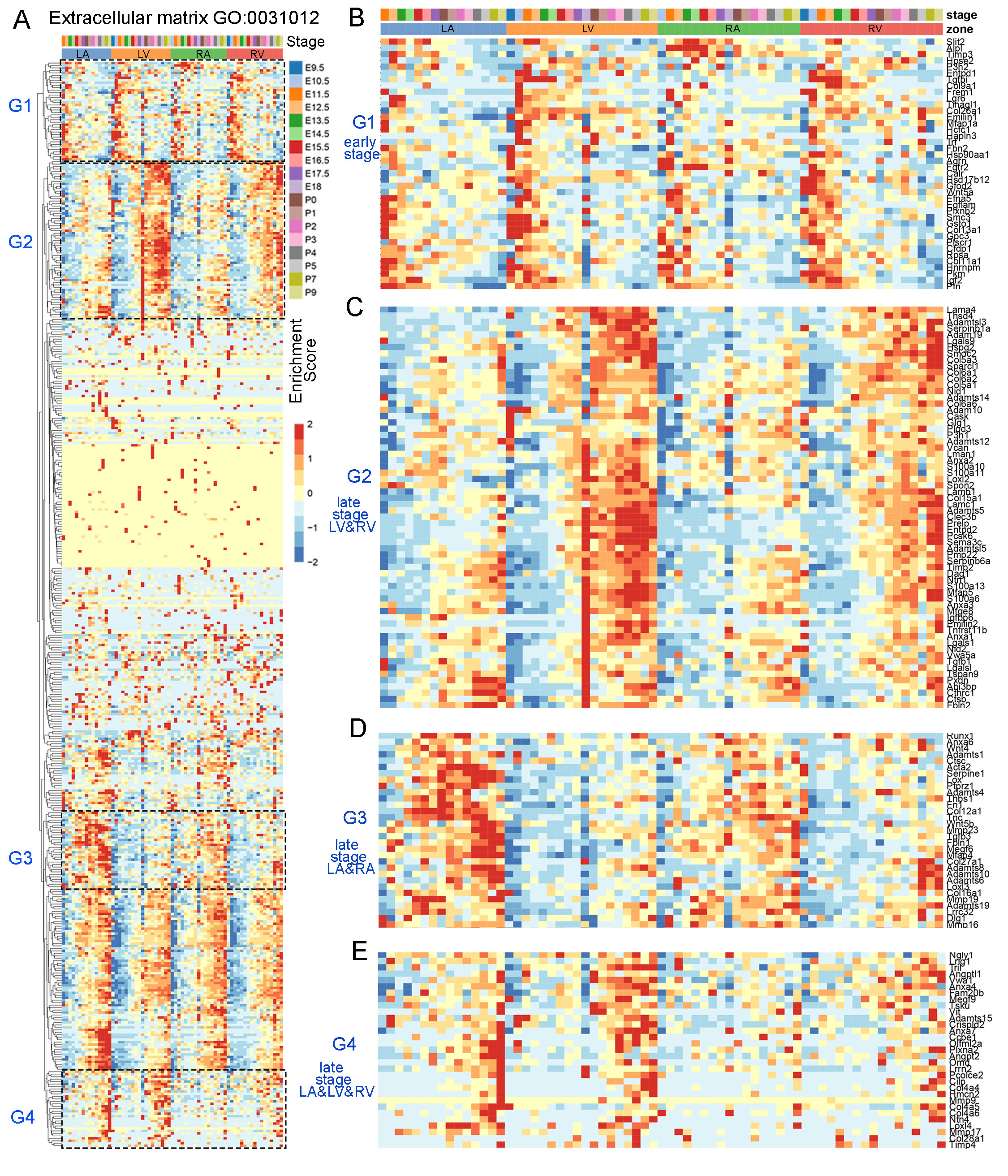

### Supplemental data 6

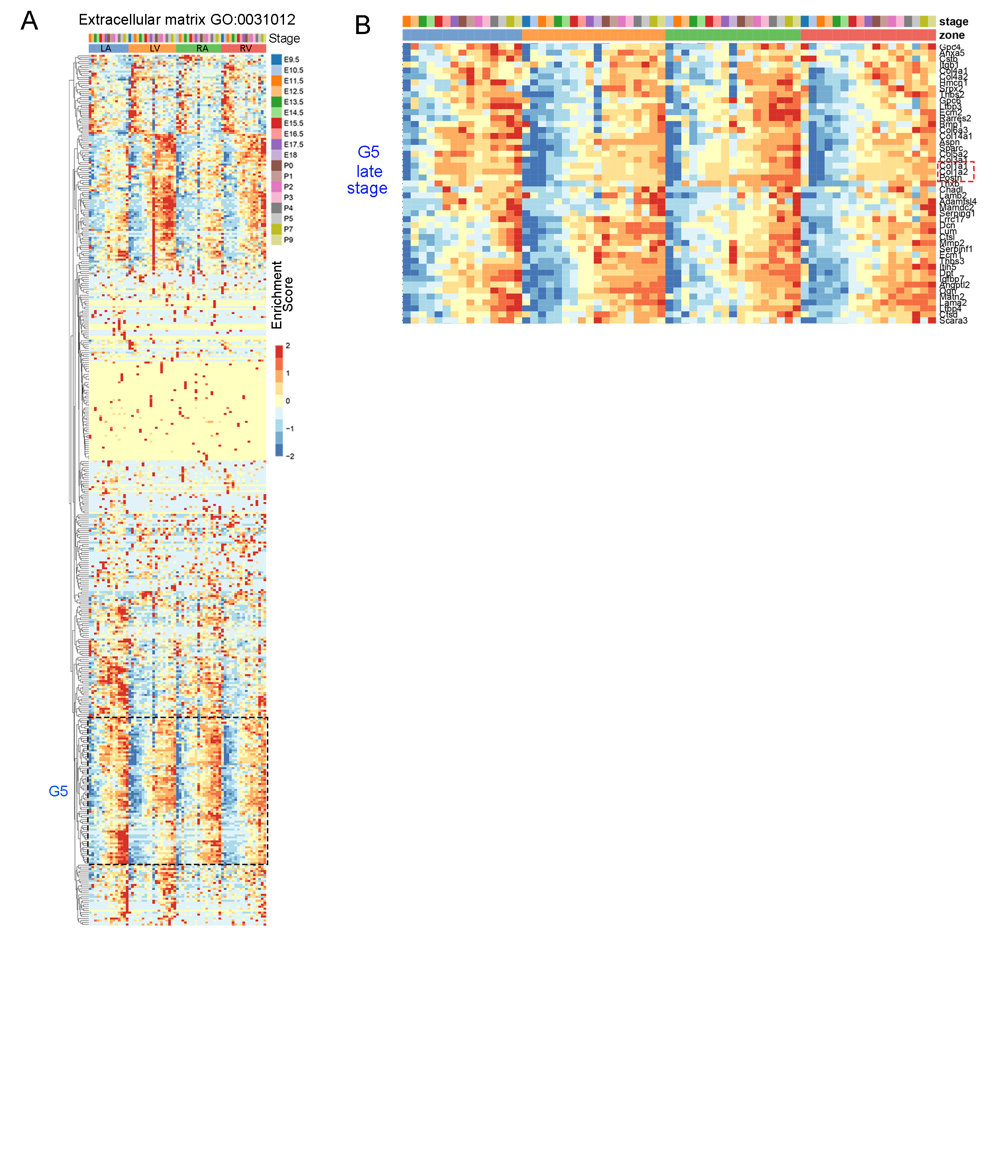

### Supplemental data 7

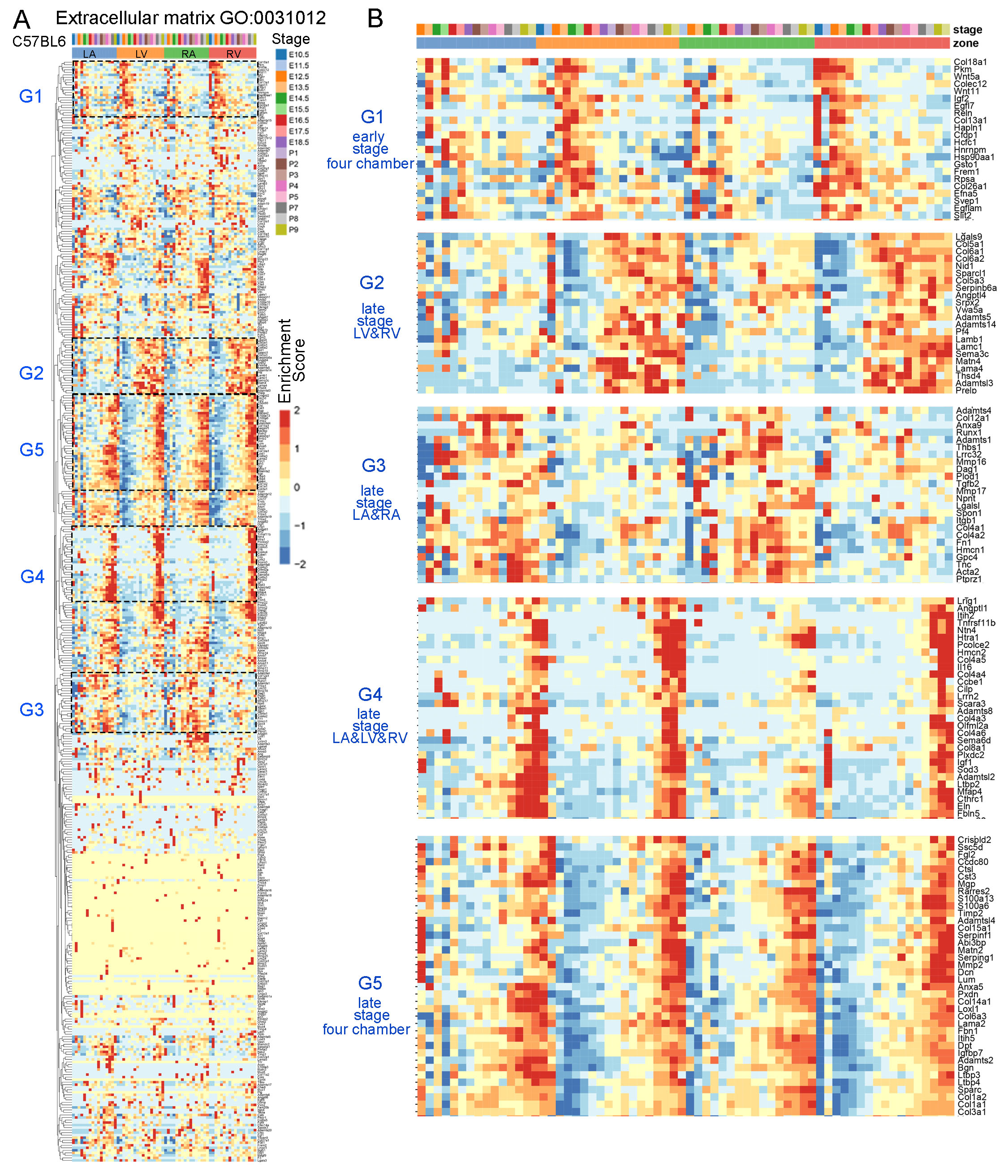

### Supplemental data 8

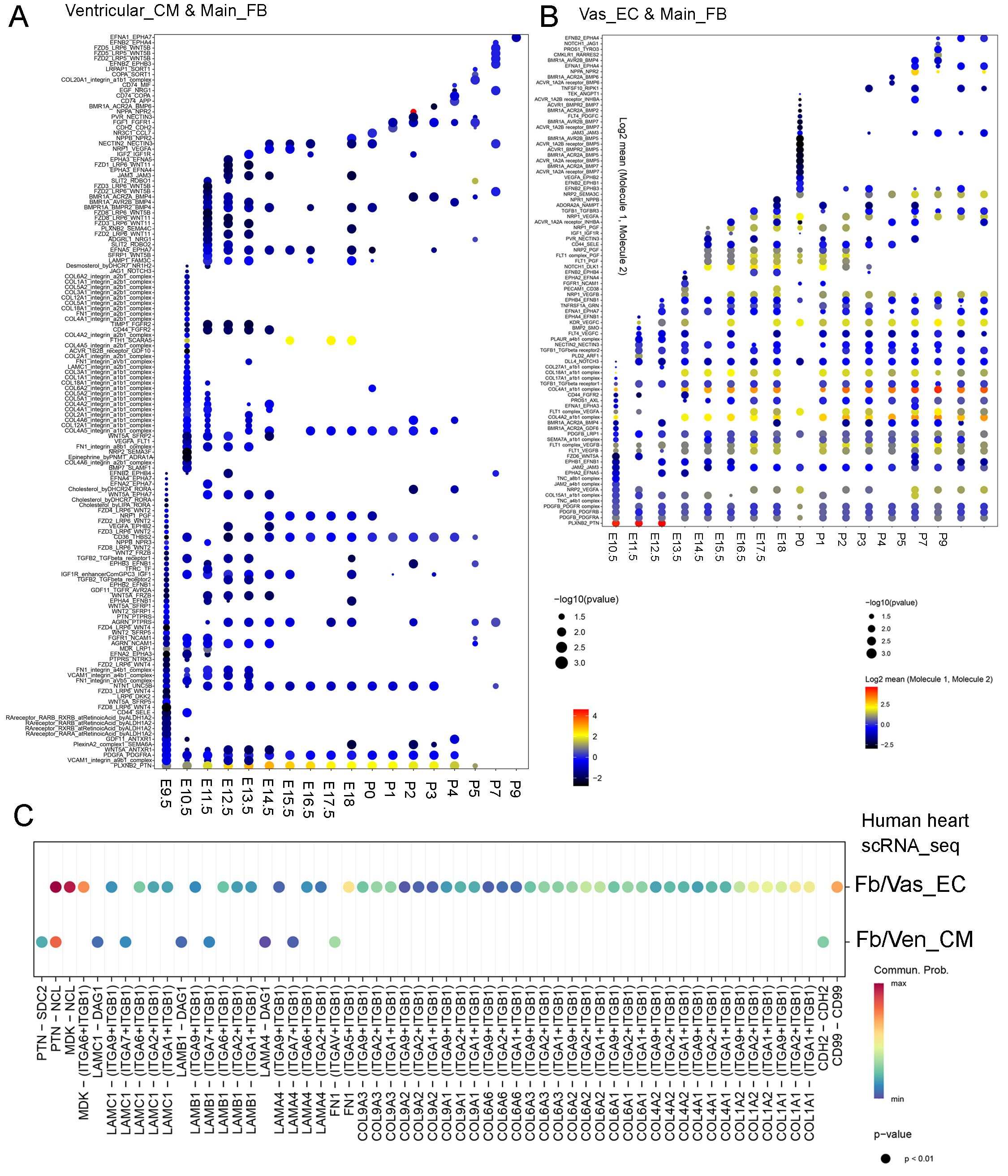

### Supplemental data 9

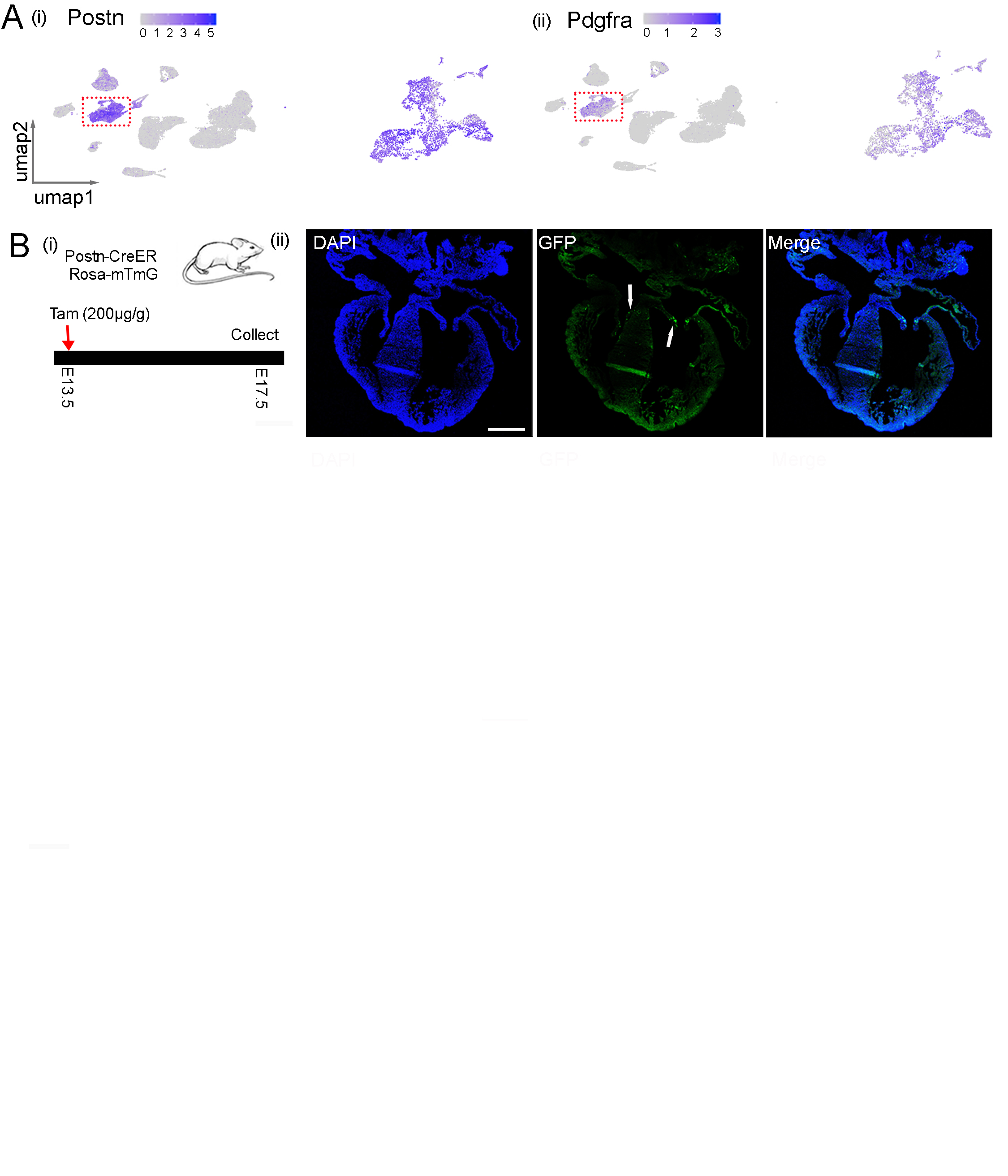

### Supplemental data 10

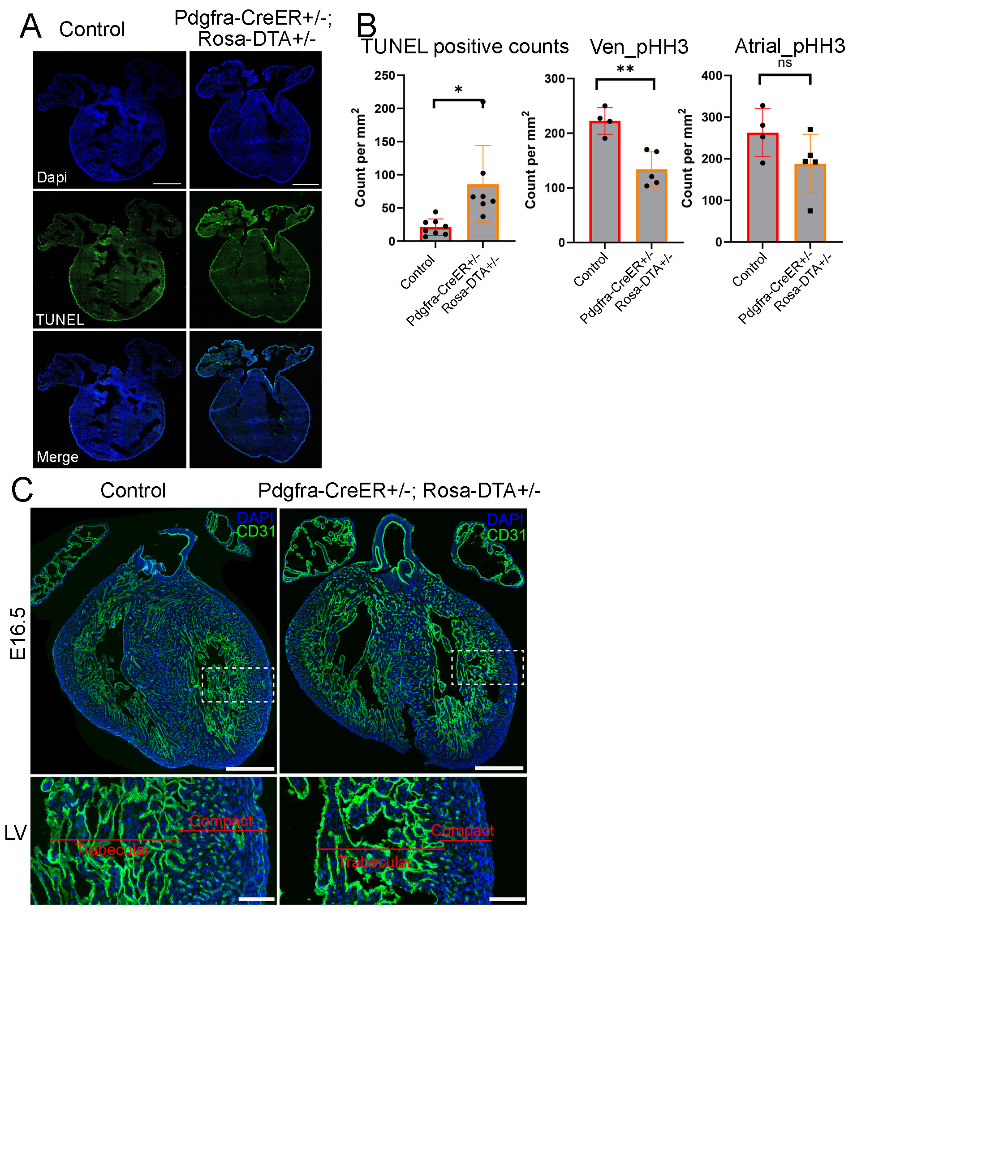

### Supplemental data 11

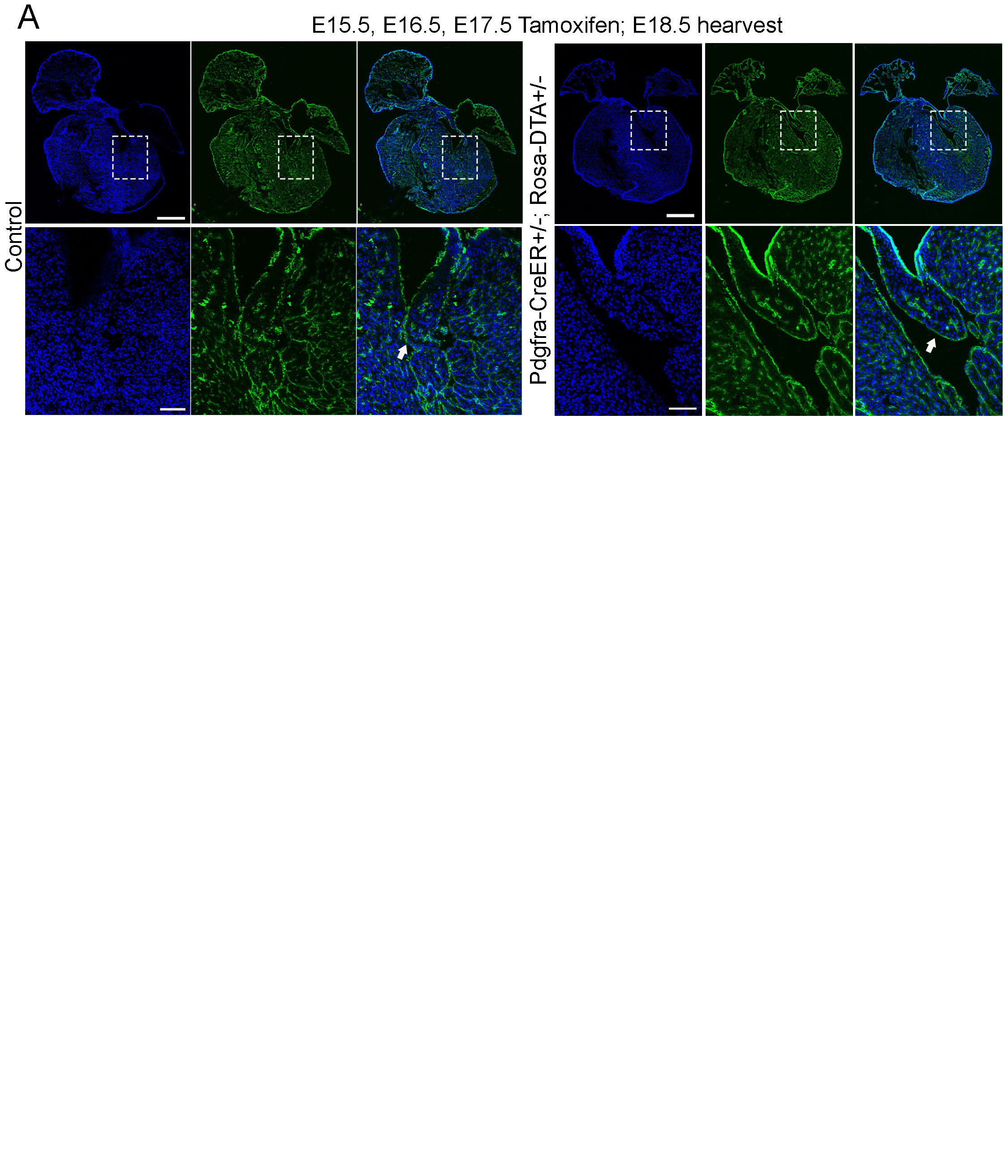

### Supplemental data 12

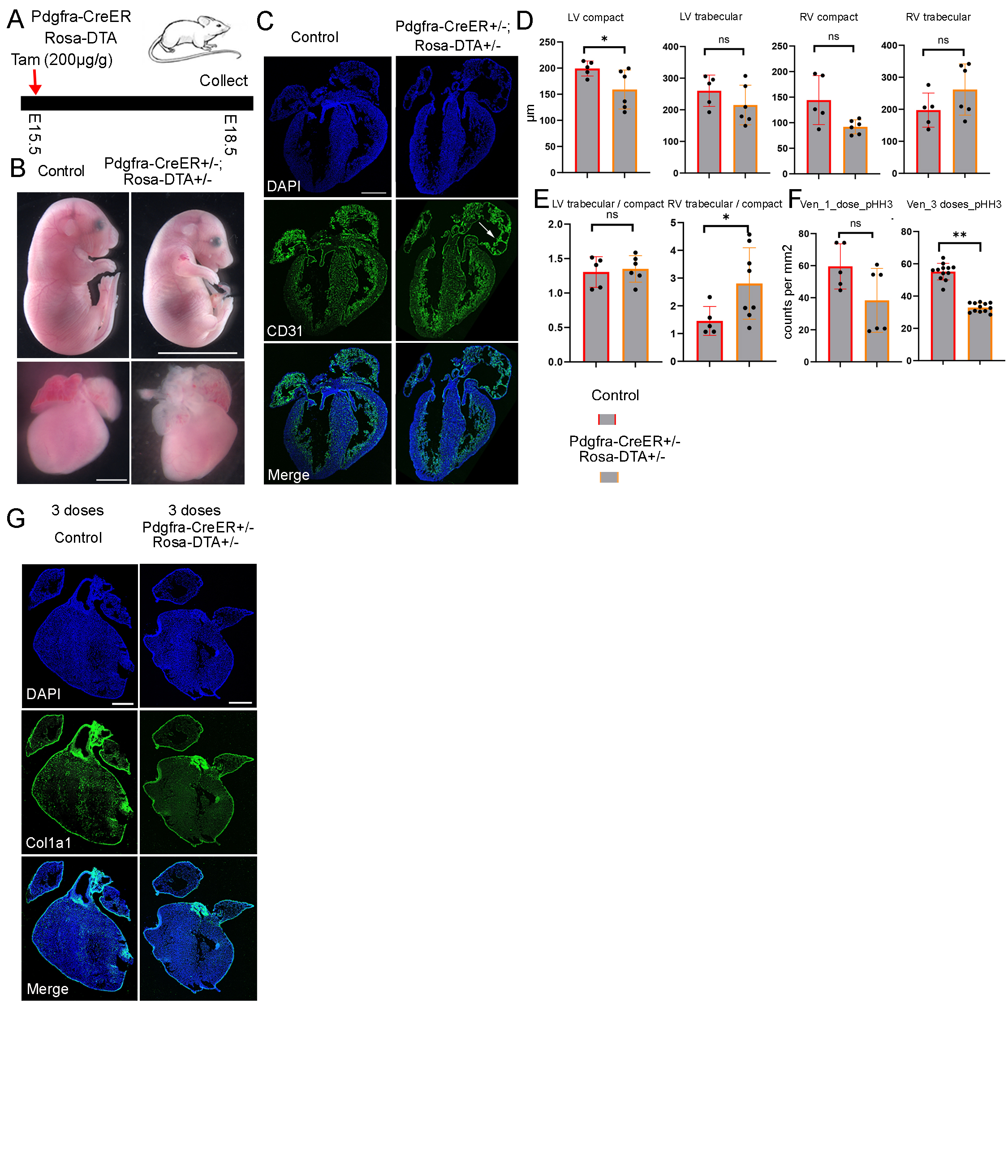

### Supplemental data 13

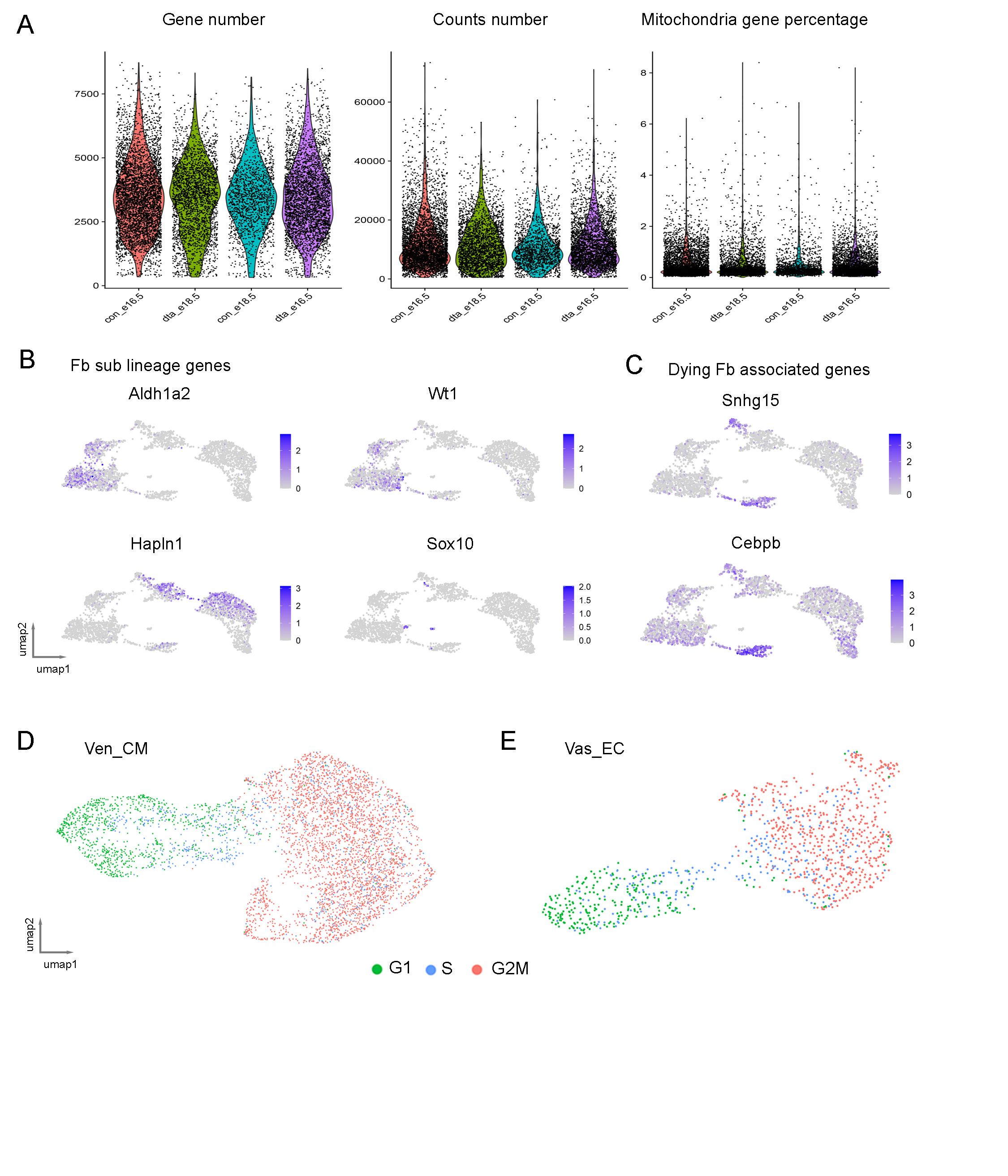

### Supplemental data 14

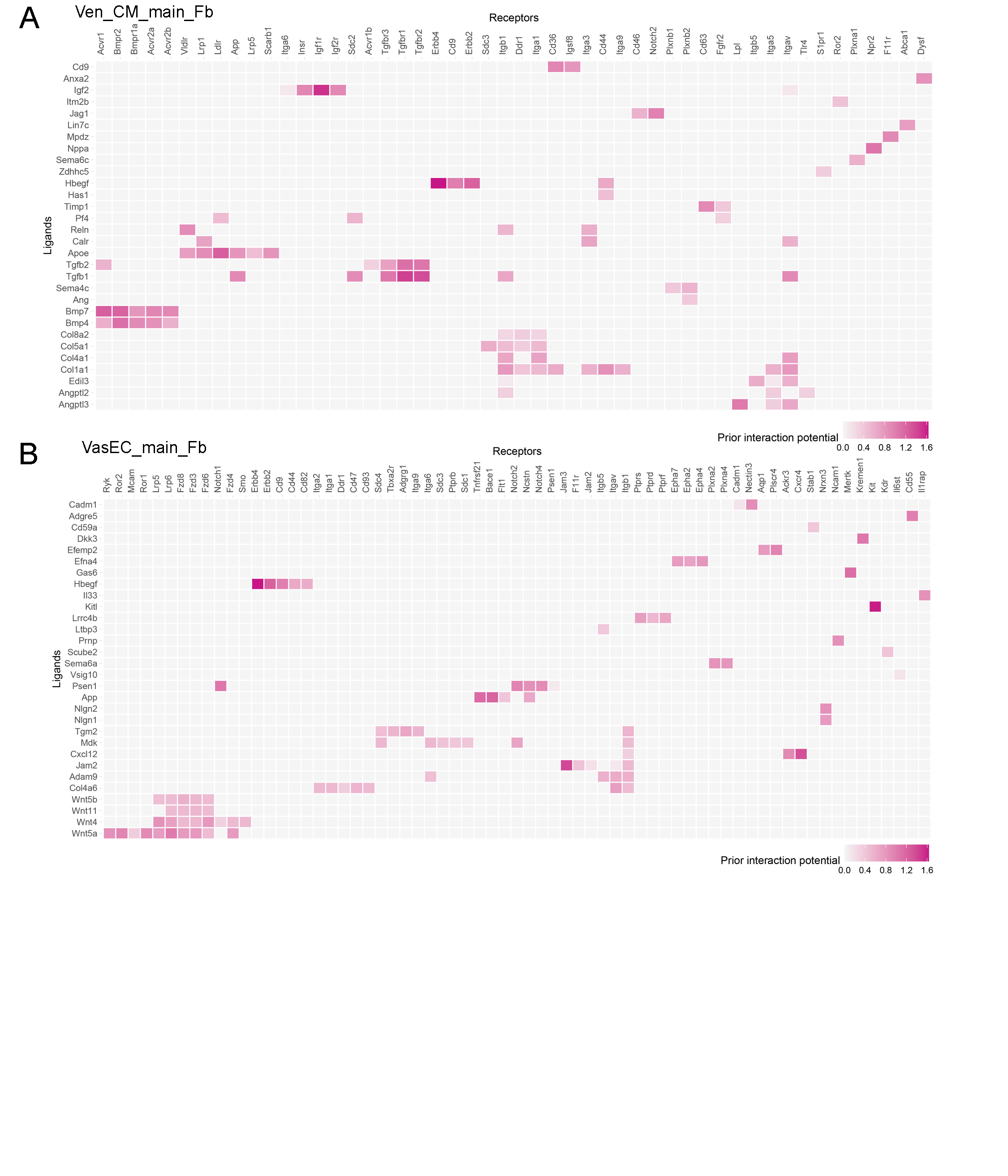

### Supplemental data 15

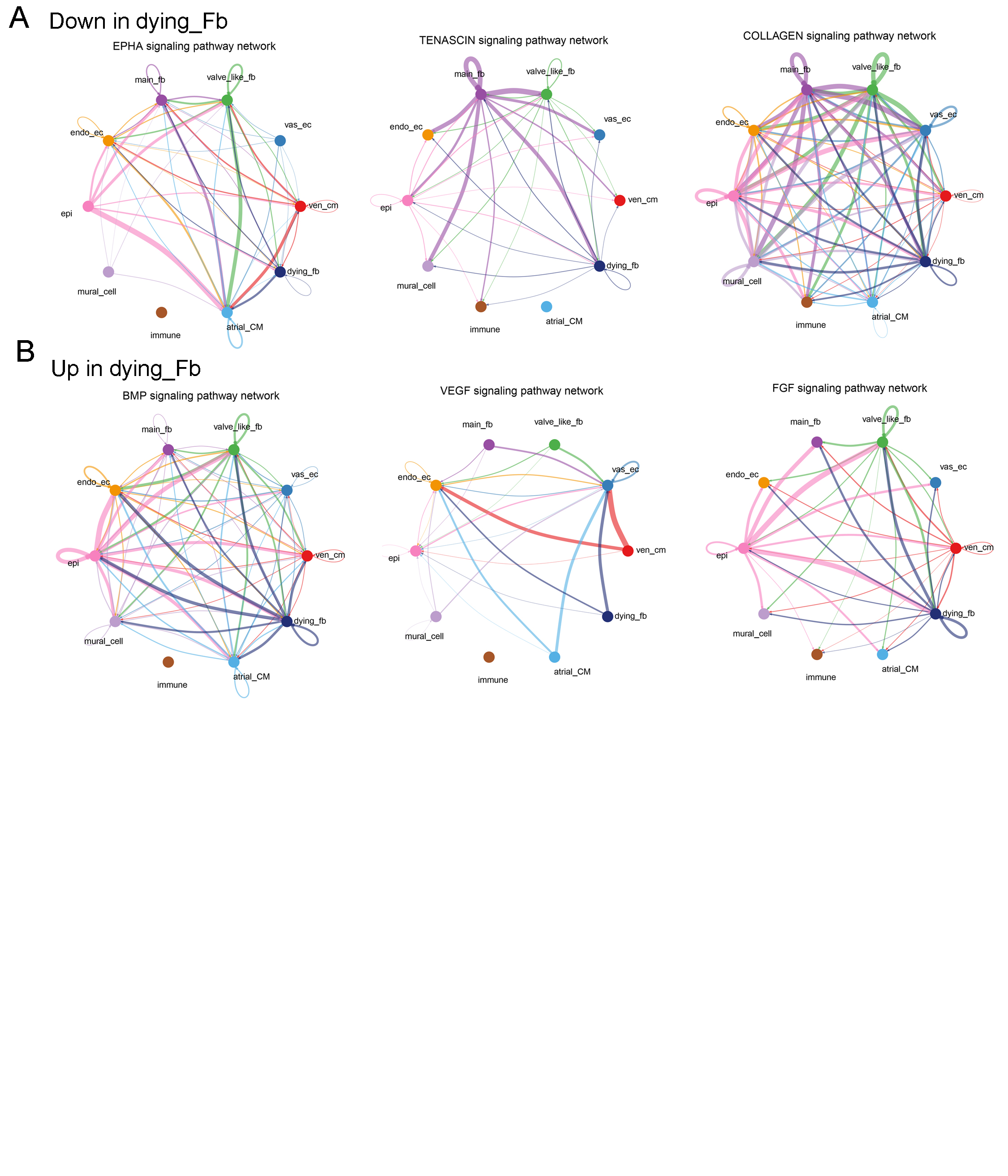

### Supplemental data 16

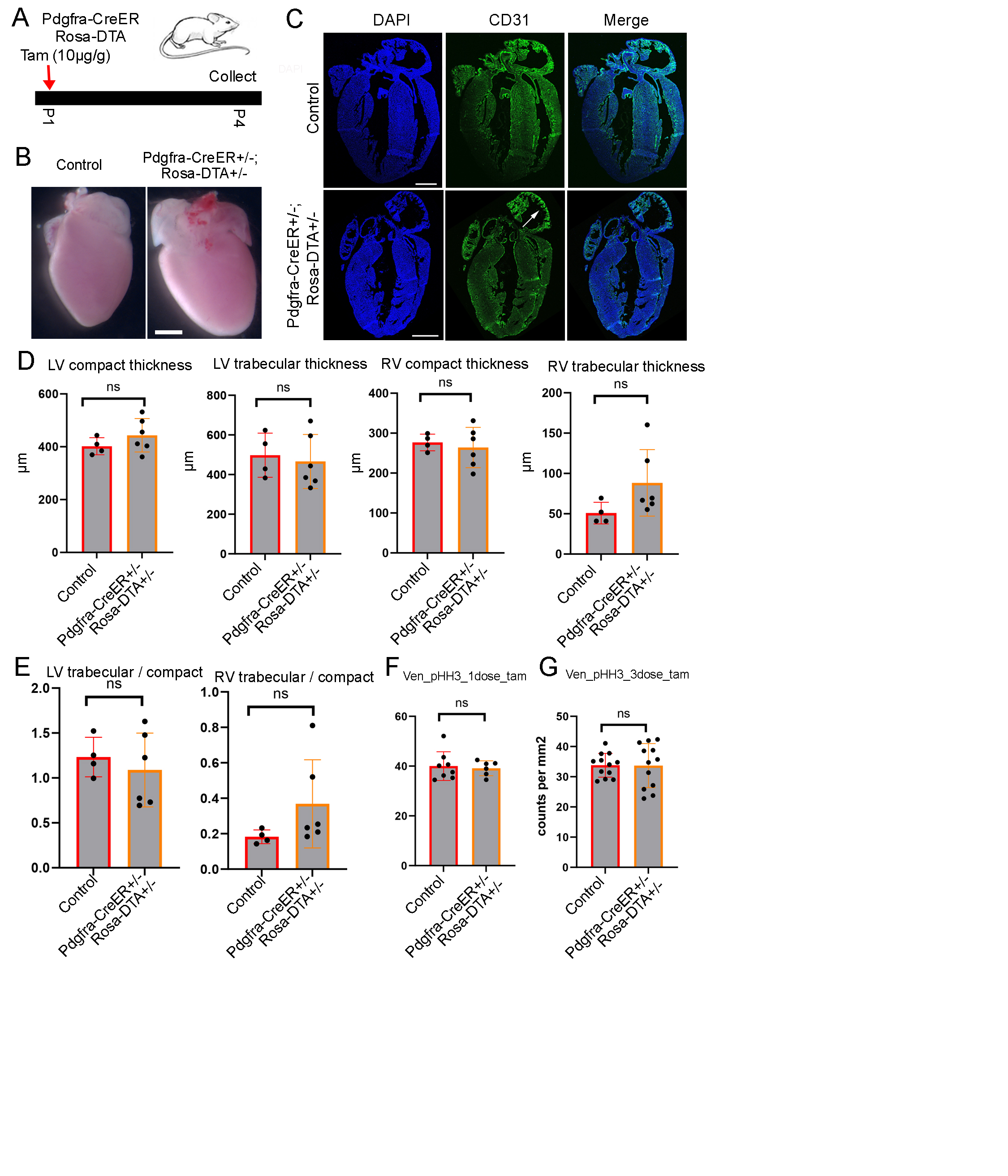

### Supplemental data 17

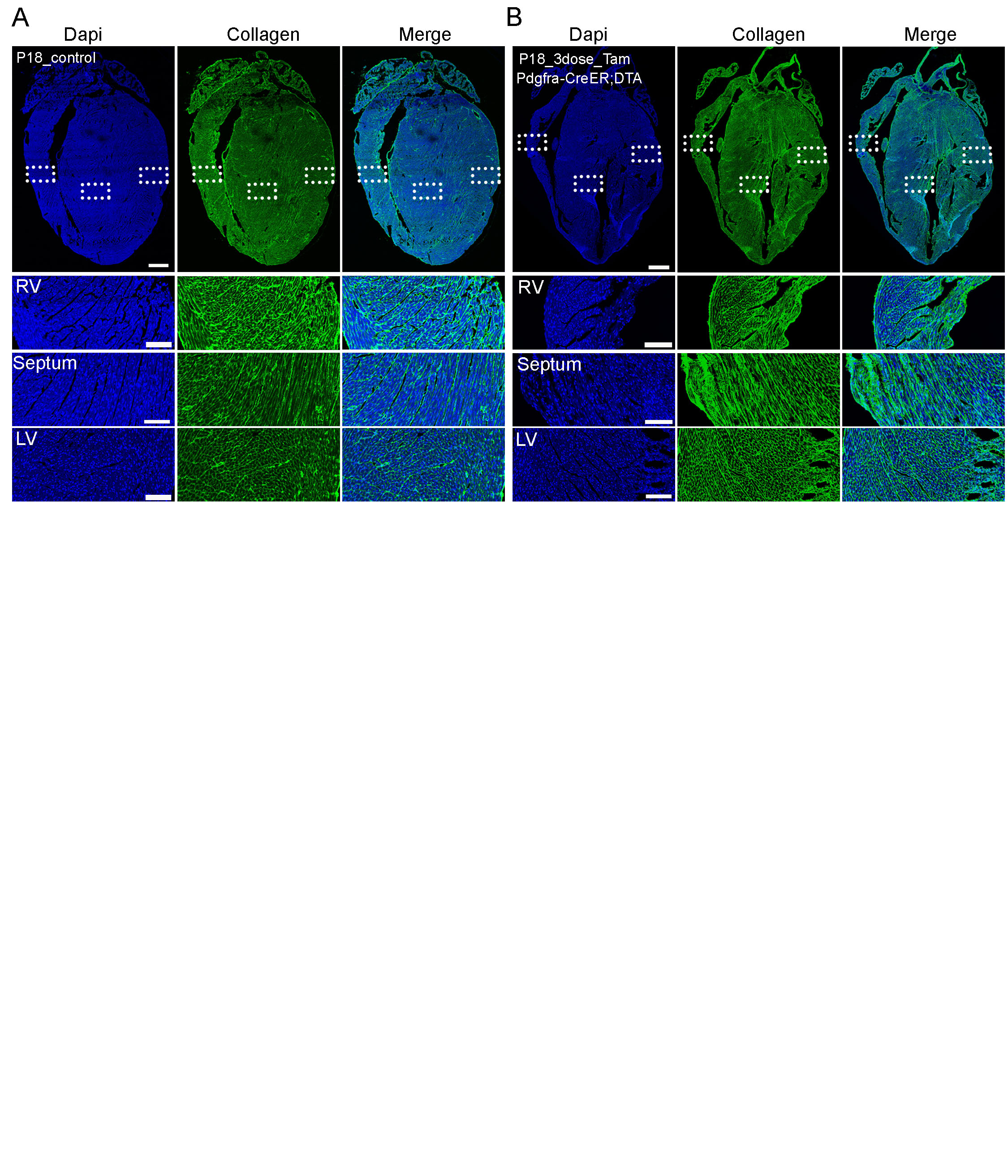

### Supplemental data 18

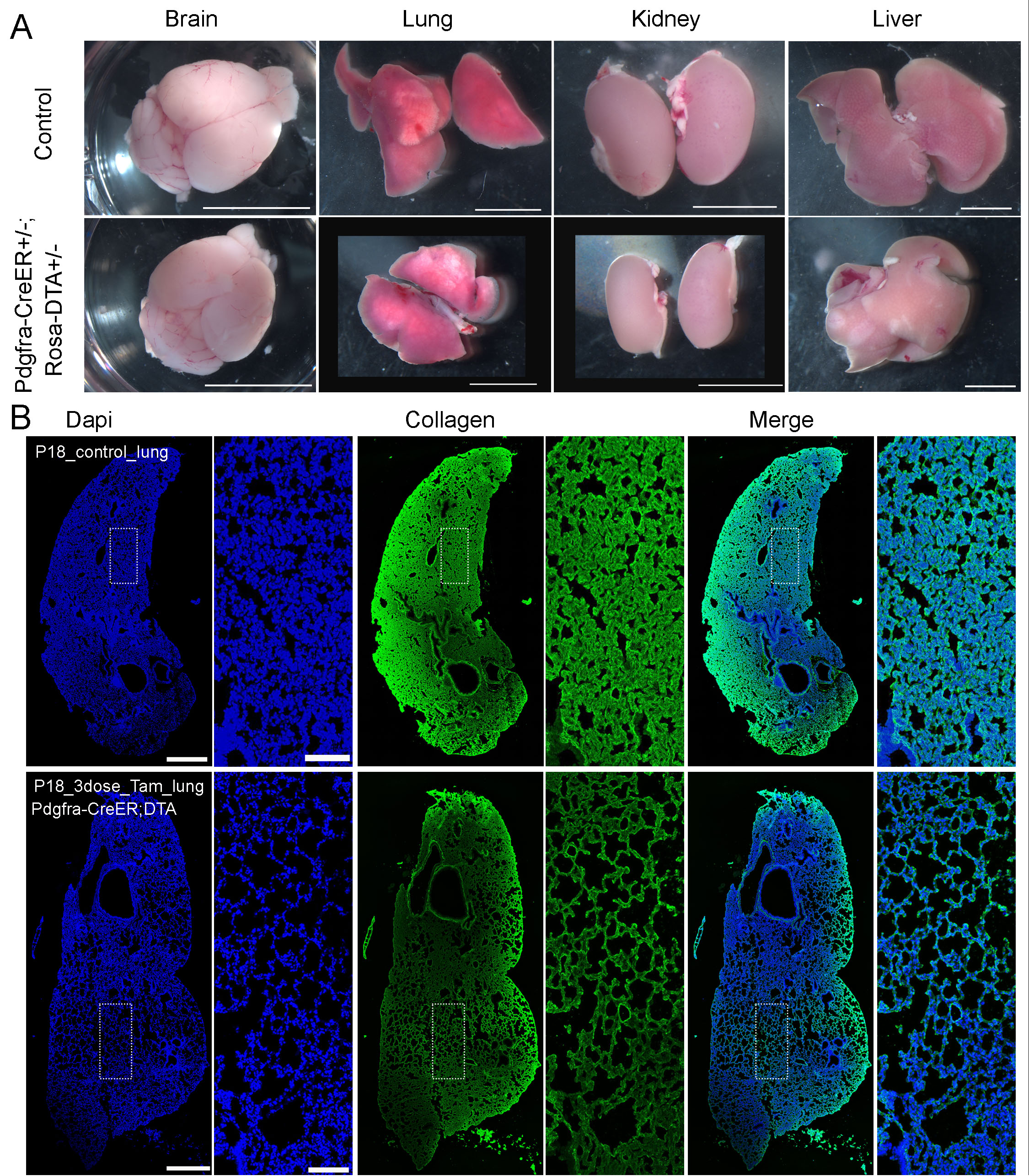
