## Supplemental table 1 for "Cardiac Fibroblasts regulate myocardium and coronary vasculature development via the collagen signaling pathway"

| **Name** | **Sequence** |
| --- | --- |
| cy5-mCol1a1-Right-1 | CCACCCCTTCACAGAGATGTTTATACGTCGAGTTGAACGTCGTAACA |
| cy5-mCol1a1-left-1 | TAGCGCTAACAACTTACGTCGTTATGAGCACCTTTGATACCAAACT |
| cy5-mCol1a1-Right-2 | ACACAATTGCACTGAGGAATTTATACGTCGAGTTGAACGTCGTAACA |
| cy5-mCol1a1-left-2 | TAGCGCTAACAACTTACGTCGTTATGAGAACGGTCTCTCCCACCCA |
| cy5-mCol1a1-Right-3 | CATGGAGATGCCAGATGGTTTTATACGTCGAGTTGAACGTCGTAACA |
| cy5-mCol1a1-left-3 | TAGCGCTAACAACTTACGTCGTTATGAGGTTCCTTCAACAGTCCAA |
| cy5-mCol1a1-Right-4 | GACTTATACCCACATAGGTCTTATACGTCGAGTTGAACGTCGTAACA |
| cy5-mCol1a1-left-4 | TAGCGCTAACAACTTACGTCGTTATGTTCAAGCAAGAGGACCAAGC |
| cy5-mCol1a1-Right-5 | GCCCCAAGTTCCGGTGTGACTTATACGTCGAGTTGAACGTCGTAACA |
| cy5-mCol1a1-left-5 | TAGCGCTAACAACTTACGTCGTTATGTCGTGCAGCCGTCCACAAGG |
